# Successive cultivation under drought selects for specific microbiome members in the wheat rhizosphere

**DOI:** 10.1101/2025.09.18.677050

**Authors:** Adele Pioppi, Sofia I. F. Gomes, Mette Nicolaisen, Xinming Xu, Ákos T. Kovács

## Abstract

Growing knowledge on plant microbiomes demonstrates the contribution of the host plant during microbiome assembly, especially under stress conditions commonly threatening crops. To dissect the influence of a plant on its microbiome, repeated cycling of microbiomes can be utilized to enhance functional properties in the enriched microbial communities. We used such a successive cultivation approach for wheat (*Triticum aestivum*) microbiome under drought conditions and selected lineages for drought resilience and susceptibility, with and without enriching the starting community with a wheat isolate library. Significant differences in the rhizosphere microbiome between selection regimes were confirmed through 16S rRNA gene amplicon sequencing. Notably, replicate lineages of each selection regime showed convergence to similar microbiomes. Specific genera were abundant depending on the selection regimes; *Stenotrophomonas* under drought resilience, while *Rahnella* under drought conditions when the strain library was added initially. Applying *Stenotrophomonas* or *Rahnella* as single inoculum did not improve drought resilience in wheat. We hypothesize that complex microbiome dynamics take place during successive cultivation, which underscores the importance of considering complex plant-microbiome systems for studying plant stress resilience. Successive cultivation remains a valuable approach for observing rhizosphere microbiome changes under different conditions.

## Introduction

Crops cultivated worldwide regularly face unfavorable environmental conditions, intensified by recent climate change, which can severely impair harvest yields, and thereby danger food security. In particular, drought has become one of the most pressing concerns that threatens natural ecosystems and agriculture, and causes major crop losses globally, while it is prospected to become even more severe in the future (Lesk, Rowhani and Ramankutty 2016). In order to maintain sufficient crop production under widespread environmental stressors including drought, there is an increasing need for complementary sustainable solutions centered around plant health and resilience (Jansson 2023).

The rhizosphere microbiome has increasingly been recognized as a key factor contributing to plant health and stress resilience (Compant *et al*. 2025). Plants interact with the microbiome in the soil to form unique rhizosphere communities which tend to differ depending on environmental conditions (Berendsen, Pieterse and Bakker 2012). Intriguingly, plants under environmental stress often shape their rhizosphere microbiomes differently than unstressed healthy plants (Sheik *et al*. 2011). Plant-microbiome interactions are especially tight during the early development of plants, and seed-borne microorganisms play an active role in mediating the plant’s interaction with the environment (Torres-Cortés *et al*. 2018). Exploiting the variability of microbial communities and their tight associations with their host is the basis of emerging microbiome engineering approaches (Berruto and Demirer 2024).

To study how microbiome assemblies are shaped by the host plant, a successive cultivation approach can be utilized. Successive cultivation generally consists of repeated cycling of the rhizosphere microbiome extracted from plants across multiple cycles of cultivation under a defined condition, based on the premise that this will produce significant shifts in microbiome composition and functionality (Swenson, Wilson and Elias 2000). This approach relies on the host plant’s interactions with its microbiome to select functionally characteristic microbial communities under a given condition. Plants themselves are not propagated through multiple generations, as the focus lies on microbiome development and not on plant adaptation. For example, successive cultivation has been shown to amplify microbiome differences between distinct plant varieties (Cordovez *et al*. 2021). Phenotypic traits of interest are typically monitored during successive cultivation and can be used to inform the selection of specific plants to be used for microbiome re-inoculation (Swenson, Wilson and Elias 2000; Panke-Buisse *et al*. 2015). This method can inform about measurable effects in the relevant plant parameters at the end of the process. Mueller *et al*. (2021) developed a microbiome conferring salt tolerance to *Brachypodium distachyon* after nine rounds of microbiome cycling under sodium salt stress (Mueller *et al*. 2021). De Zutter *et al*. (2021) performed consecutive microbiome cycling in maize under low phosphorus availability thereby achieving the enrichment of phosphorus-solubilizing bacteria in the rhizosphere microbiome and an improved P status in the plant (De Zutter *et al*. 2021).

In the current literature, there are several examples of successive cultivation being employed to obtain functionally beneficial microbiomes, however there is limited investigation on exactly which microbiome components mediate the effects. Furthermore, microbiome cycling is typically done by choosing the best performing plants, without further knowledge or emphasis on microbiomes associated with poorly performing plants. Microbiomes associated with plants showing less-preferred phenotypes are often inspected to identify key groups enriched under a stress condition, such as under active fungal infection (Li *et al*. 2021b). However, this aspect is neglected in successive cultivation experiments.

Here, we performed successive cultivation of wheat rhizosphere microbiome under drought conditions, including separate plant selection lines – hereafter called lineages – for resilience and susceptibility to drought, as well as lineages of random plant selection using watered conditions. Moreover, we enriched the initial endogenous seed-borne community with a bacterial library of wheat isolates. We hypothesized that this would result in more diverse microbiomes compared with those formed solely by cycling seed-borne bacteria. Because all individual plants are assumed to share the same genetic makeup, the selection of plants in this experiment targets their phenotype as a product of plant-microbiome interactions. We aim to (i) apply successive cultivation to wheat under drought conditions and observe the microbiome divergence between healthy and stressed plants; (ii) identify which components of the wheat rhizosphere microbiome distinguish more resilient plants from more susceptible plants during drought stress.

## Materials and methods

### Bacterial library

A bacterial library was prepared for plant inoculation that consists of 84 strains isolated from wheat rhizosphere and rhizoplane (Ahmad *et al*. 2022) and two *Bacillus subtilis* strains isolated from wheat rhizoplane samples using heat treatment and selecting for structured colonies (Kiesewalter *et al*. 2021; Xu *et al*. 2024), for a total of 86 strains (Fig. S1). The strains were identified by sequencing of the 16S rRNA genes using primers 27F 5’-AGAGTTTGATCMTGGCTCAG-3’ and 1492R 5’-TACGGYTACCTTGTTACGACTT-3’ at Eurofins and taxonomy was identified using NCBI Blast (see Table S1 depicting the strain list and taxonomic assignment and Dataset S1 for sequences). All strains were grown at 28°C for 48h and mixed equally after adjusting each strain to an optical density at 600 nm (OD_600_) of 0.1 to prepare the inoculum for the experiment.

### Successive cultivation setup and treatments

Peat free pellets were used as substrate for wheat cultivation. First, the total water holding capacity (WHC) of the pellets was determined (Margesin and Schinner 2005), and expected soil weight was calculated on a step scale from 10% WHC to 70% WHC. Before each cultivation cycle, new dry pellets were hydrated with 25 mL 0.5% Hoagland’s solution each, then autoclaved two days before planting (d-2), and again on the day of planting (d0), after which they were assumed to be at 70% of WHC. The total weight of pot and pellet was adjusted to 25g before planting.

Winter wheat seeds (Sheriff variety; Sejet Plant Breeding, Horsens, Denmark) were surface sterilized by shaking with 2% sodium hypochlorite for 20 min, then washed 5 times with MilliQ. The seeds were germinated on filter paper soaked with MilliQ for 2 days in sealed petri dishes kept in the dark. One seedling was sown in each pot (3.5 × 3.5 × 4.0 cm). Plants were grown in a growth cabinet (Binder, Tuttlingen, Germany) with 16h of light at 22°C and 8h of darkness at 18°C. Each cultivation cycle lasted 16 days in the growth cabinet. On the last day (d16) of each cultivation cycle, one plant was selected from each selection lineage in order to re-inoculate its rhizosphere extract into the following cycle of plants on their respective d2. Three different plant selection rationales were used: (i) random selection for plants under watered conditions, (ii) selection of the most resilient plant under drought conditions, and (iii) selection of the most susceptible plant under drought conditions (Fig. 1). Resilient plants were visually identified by having a majority of upright fresh leaves, while susceptible plants were visually identified by a majority of wilting and dry leaves. Each selection rationale was applied to three replicate lineages which received library inoculation at the beginning of cycle 1, as well as three replicate lineages which were not inoculated with the library. Each lineage included five plant replicates in each cycle.

**Figure 1.**
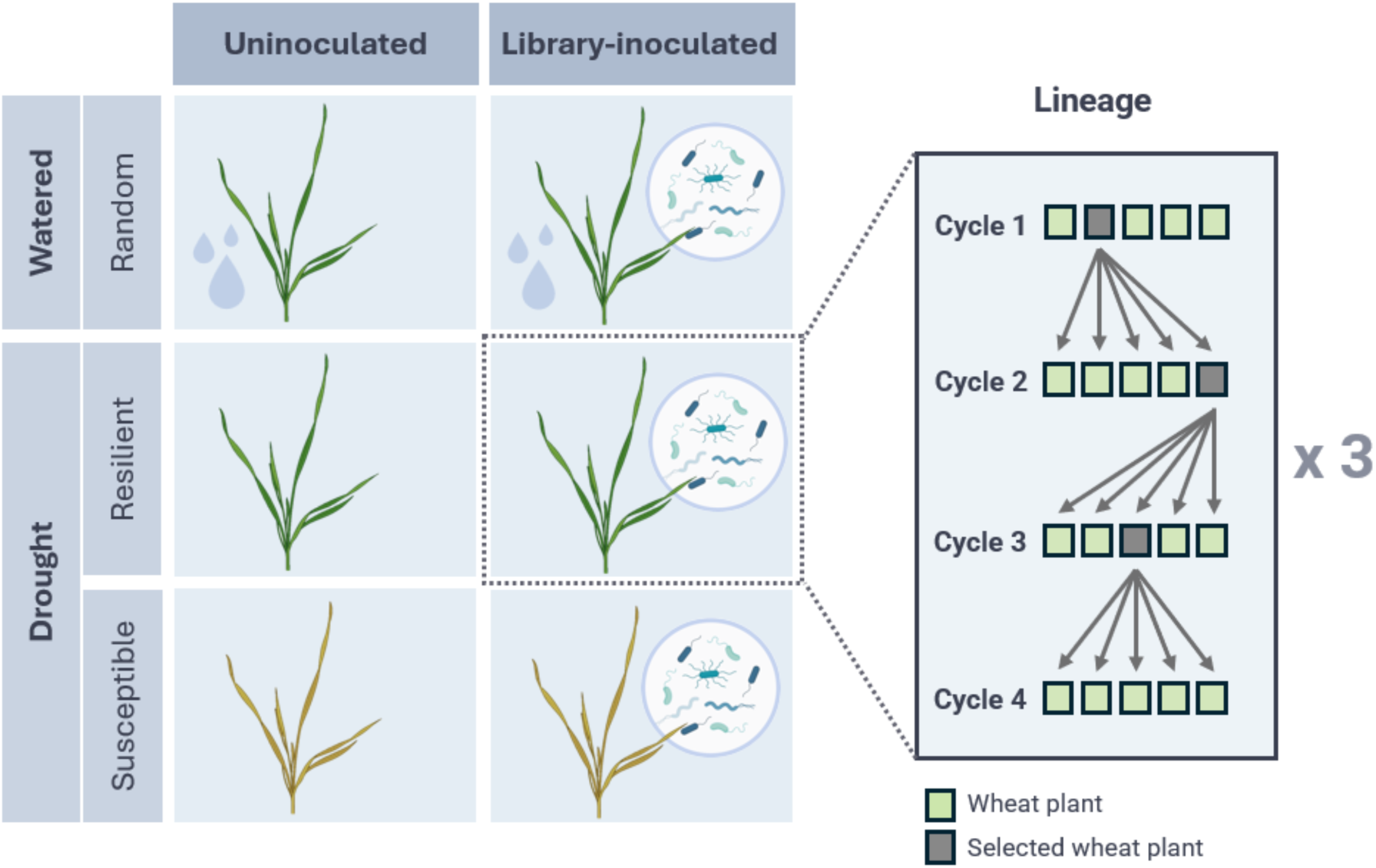
Schematic representation of the inoculation treatment and selection regimes used in the successive cultivation setup. Watered plants were selected randomly, while plants under drought were selected either for drought resilience or susceptibility. Each combination included three replicate lineages with three cycles of rhizosphere re-inoculations from a selected plant from cycle 1 to cycle 4. The depicted lineage scheme depicts an example for plant selection and reinoculations between cycles.

### Watering and application of drought stress

Plants were watered with autoclaved tap water based on daily measurements of total weight of the pot. Pots in drought treatments were watered to the target weight (corresponding to 70% WHC) until d5, then allowed to progressively dry out to approximately 10% WHC with no further watering until d16. Pots in watered treatments were watered consistently to keep 60-70% WHC. The placement of replicate pots was regularly shifted to ensure even drying.

### Cultivation cycles and re-inoculation workflow

The successive cultivation consisted of four plant growth cycles. In each cycle, wheat seedlings were planted on d0 and inoculated on d2 as follows. On d2 of the first cycle 2mL of the mixed library were inoculated at the base of each plant, while in following cycles the rhizosphere extract from a selected plant in the previous cycle was used for inoculation. Rhizosphere extraction from the selected plants was done on d16 of each cycle following the approach explained in Cordovez et al. (2021) with modifications (Cordovez *et al*. 2021). Briefly, bulk soil was gently removed, and the roots with remaining rhizosphere soil were cut into a 50 mL falcon tube containing 7 mL of 10 mmol/l MgSO_4_ (Roth) buffer. The tubes were vortexed at high speed for 1 min, then treated in a water bath sonicator for 2 min, and finally vortexed again for 30 s. The solution was filtered through 2 layers of sterile Miracloth (Millipore), and 2 mL of it was administered to each plant of the new cycle on d2. On d16 of the 4^th^ and last cycle (Fig. S2), rhizosphere samples were extracted from all plants as described up to the filtering step. The liquid extract was stored at -20°C for further use.

### DNA extraction and 16S rRNA gene amplicon sequencing

DNA extraction from the rhizosphere samples was done with DNeasy PowerSoil Pro Kit (QIAGEN). The V3-V4 region of the 16S rRNA gene was amplified with barcoded primers 5’-CCTACGGGNGGCWGCAG-3’ and 5’-GACTACHVGGGTATCTAATCC-3’. A 50 µL PCR reaction contained 17.8 µL MilliQ, 25 µL of TEMPase Hot Start 2× Master Mix (AMPLIQON), 1.6 µL of each primer (10 µmol/l), and 4 µL DNA template. The PCR program consisted of an initial denaturation step at 95°C for 15 min, followed by 30 cycles of 95°C for 30 s, 62°C for 30 s, and 72°C for 30 s, with a final extension at 72°C for 5 min. The PCR products were purified with the NucleoSpin Gel and PCR clean-up kit (Macherey Nagel) according to the manufacturer’s specifications, and finally pooled together in equimolar ratios. The samples were sequenced by NovaSeq PE250 with 2G raw data per sample at Novogene Europe Company Limited.

### Bioinformatics and analysis of community composition

A total of 2.3 million high-quality reads were obtained, corresponding to an average of 63,727 reads per sample. Amplicon sequence data were analyzed in R. Cutadapt was used to remove barcoded primer sequences. DADA2 was then used to filter, trim, and denoise reads, followed by merging paired-end reads (Callahan *et al*. 2016). Finally, chimeras were removed, as well as non-bacterial sequences, chloroplast and mitochondrial sequences. Amplicons were compared to the NCBI nucleotide (nt) database using BLASTN (max_target_seqs = 3), and the top hits were used for taxonomic classification. Amplicon counts were rarefied to an even sequencing depth using the rarefy_even_depth function in the phyloseq R package (McMurdie & Holmes, 2013), with random subsampling without replacement. The R package ampvis2 was used to analyze the community composition and diversity (Andersen *et al*. 2018).

### Testing specific strains from the wheat isolate library under drought conditions

Based on our results highlighting differentially enriched bacterial groups between treatments, we proceeded to test isolates belonging to two genera in a follow-up cultivation experiment. One *Rahnella* strain and 10 *Stenotrophomonas* strains were tested in a single cultivation cycle for their ability to positively influence wheat drought resilience. Preparation for cultivation was carried out as described for the successive cultivation setup, including seed sterilization, germination (lasting four days), and autoclaving and hydration of the soil pellets. Treatment with *Rahnella* included *Rahnella sp.* strain D9, while treatment with *Stenotrophomonas* included an equally distributed mix of *Stenotrophomonas* strains A7, C3, C12, D2, D3, D8, D10, E2, F5, and F12. The two inoculants were prepared from culturing the strains overnight in lysogeny broth (LB Lennox, Carl Roth) at 28°C and subsequent dilution to an optical density at 600 nm (OD_600_) of 1.0 using MilliQ water. For the *Stenotrophomonas* inoculum, the 10 strains were mixed equally before inoculation. A diluted LB solution without addition of bacteria was prepared as a control treatment. 2 mL of the inoculum was administered at the base of the planted seedlings. The cultivation lasted 24 days. Watering with autoclaved tap water was applied up to 70% WHC every day for all treatments up until d12, after which it stopped for the three treatments under drought. The health status of plants under drought was evaluated based on a preset scoring system: No wilting (1), slight leaf wilting (2), moderate wilting and drying leaf tips, (3) drying stem and severe wilting (4).

### Statistical analyses

Statistical differences between Shannon alpha-diversity of treatments were evaluated by one-way ANOVA and post-hoc Tukey HSD test based on sample richness data (Tukey 1949). Dissimilarity between sample compositions was measured by calculating Bray-Curtis distances using the vegan package (Oksanen *et al.* 2001; Dixon 2003), and were visualized with Principal Coordinate Analysis (PCoA) plots. Microbiome differences between all treatments were done with a PERMANOVA with 999 permutations using the adonis2 function. Differential Abundance Analysis (DAA) to highlight differentially enriched genera between treatments was done by the ANCOM-BC method including bias correction. The bonferroni method was used to adjust p-values. To avoid creating artifacts due to genera being very low or absent from a given treatment, the analysis focused on genera with overall relative abundance >1%. Genera with p-value=0 for a given comparison were excluded for the same reason. A q-value <0.1 was the cutoff for significance. Performance scores of different inoculation treatments for testing specific strains from the wheat isolate library under drought conditions were analyzed with a linear mixed-effects model (treatment, day, treatment × day as fixed effects; plant as a random effect), followed by Tukey-corrected pairwise comparisons.

## Results

### Rhizosphere bacterial community diversity after successive cultivation

Wheat rhizosphere microbiome was enriched in successive cultivation with selection for resilient or susceptible plants under drought, in addition to randomly selected plants grown in watered condition. The experiments were performed with initial inoculation of the 86 members strain library (denoted as library-inoculated) or without such inoculation (depicted as uninoculated). Importantly, the seed bacteria persisting after surface sterilization, including endophytes, contribute to the rhizosphere microbiomes. A total of 846 ASVs were detected in the final rhizosphere microbiomes. Plants resulting from lineages selected for drought resilience and susceptibility developed a rhizosphere microbiome with significantly lower alpha diversity (Shannon), compared to ones under watered conditions, both in library-inoculated and uninoculated treatments (Fig. 2). There were no significant differences in alpha diversity between resilient and susceptible-selected lineages.

**Figure 2.**
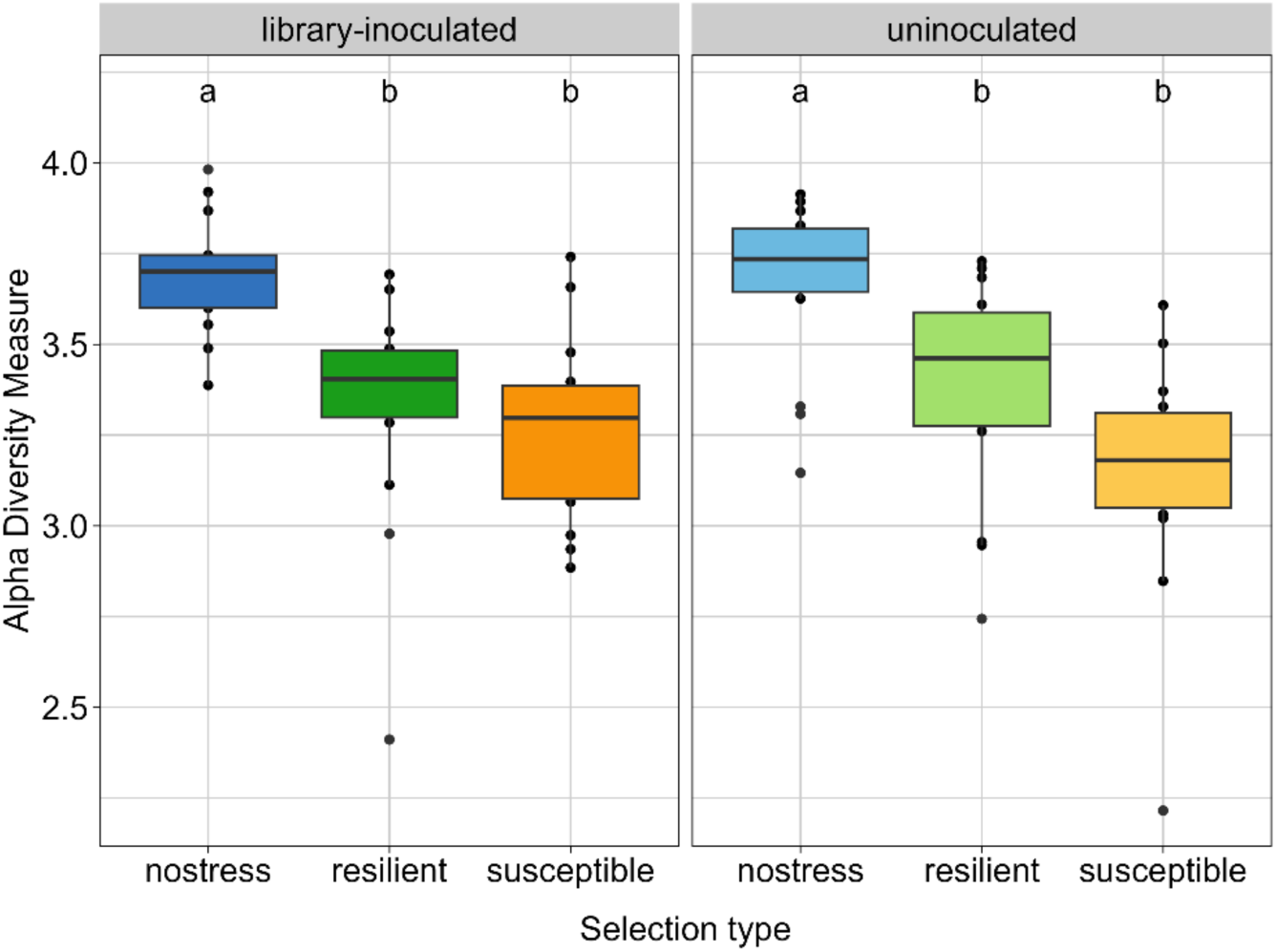
Alpha diversity (Shannon) of rhizosphere bacterial communities, distinguishing between uninoculated and library-inoculated lineages. Samples are grouped together by inoculation treatment and selection regime. Different letters above boxes indicate differences between groups. Groups not sharing a letter differ significantly.

We assayed whether the application of different selection types and watering treatments resulted in distinct rhizosphere microbiomes. The 6 treatments were distinguishable on a PCoA plot, where most sample types clustered separately (Fig. 3A). All 6 combinations of selection types and treatments were significantly different from each other (Table S2, Adonis PERMANOVA) except for the library-inoculated resilient and susceptible treatments (p adjusted=0.165). The library-inoculated treatments clustered separately from the uninoculated treatments as a whole. The inoculation factor and the selection type applied were both significant determinants of the rhizosphere microbiome (p=0.001). Taken together, these results indicate that successive selection of drought-resilient, drought-susceptible, and non-stressed plants produced different rhizosphere microbiomes, and that the initial starting community was an important factor in their development.

**Figure 3.**
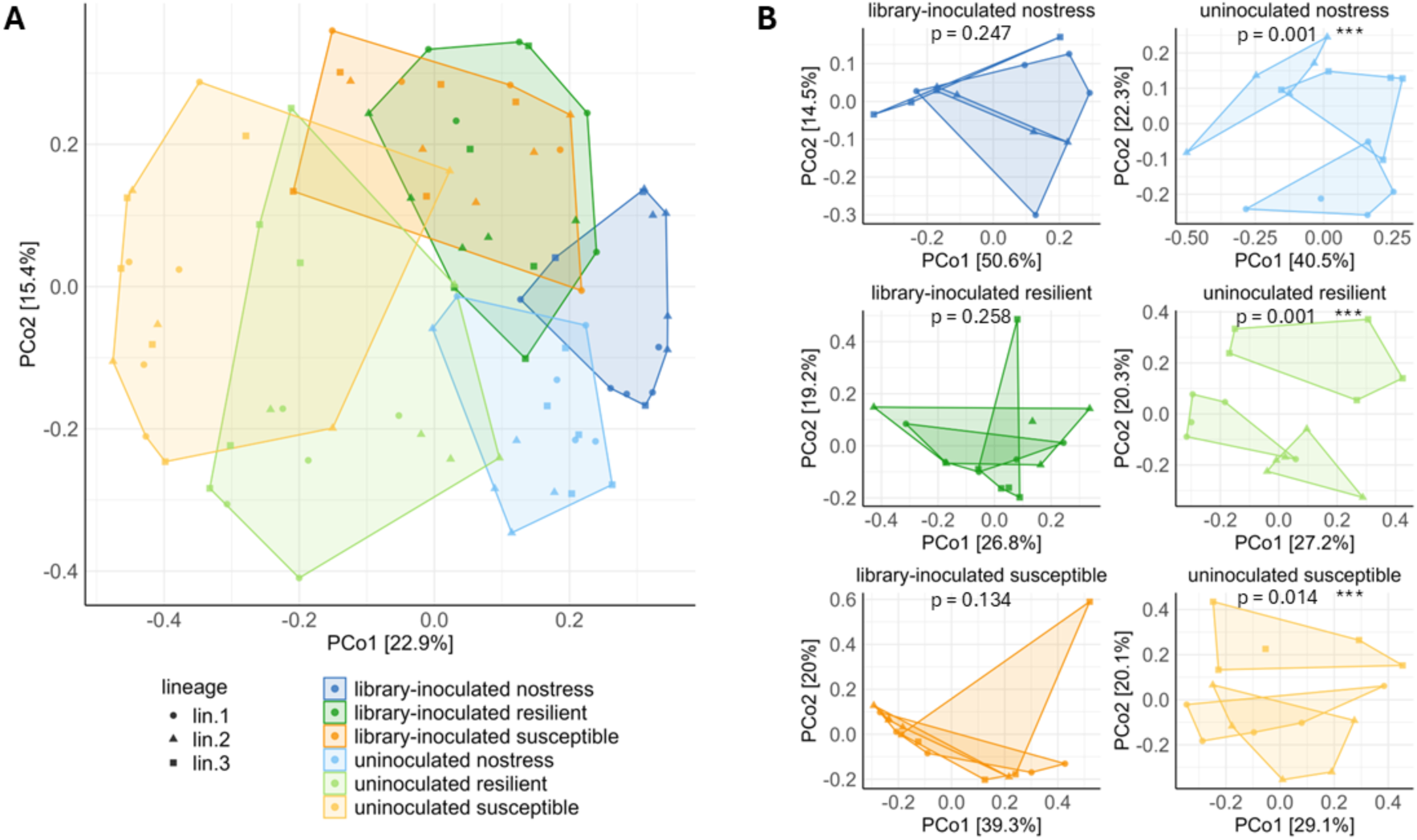
(A) PCoA plot representing diversity between selection types and inoculation treatments. ‘uninoculated’ refers to uninoculated lineages, and ‘library’ refers to library-inoculated lineages. Each sample is depicted with a data point, with a respected shape depending on the treatment condition. (B) Separate PCoA plots showing the diversity among lineage triplicates of each selection type and inoculation treatment. Significant differences between replicate lineages were tested by adonis PERMANOVA (p-value included in each plot). Significant differences between replicate lineages are depicted with an asterisk.

### Convergence and diversification of selection lineages

The successive cultivation was carried out in 3 replicate selection lineages for each of the 6 treatment combinations. We tested each treatment combination to determine whether the three lineages converged to similar microbiomes at the end of the last cultivation cycle. As determined by the PERMANOVA, the three lineages of each library-inoculated treatment do not differ significantly from each other (nostress: p=0.247, resilient: p=0.258, susceptible: p=0.134) (Fig. 3B). Instead, the three lineages of each uninoculated treatment were significantly different from each other, particularly the nostress (p=0.001) and resilient-selected lineages (p=0.001), while a higher degree of similarity was found among the susceptible-selected triplicate lineages (p=0.014).

### Microbiome composition

We examined the bacterial composition of the collected rhizosphere microbiome samples at the end of the last cultivation cycle. At least 54 genera were identified across all samples, with varying relative abundance within treatments. The 15 most abundant genera overall represent 96.37% of total read counts (Fig.4). Across treatments and selection types, a few genera emerge with higher overall relative abundance, including *Pseudomonas*, *Stenotrophomonas*, *Rahnella* and *Paenibacillus*. We note that several library genera were also found in uninoculated lineages, indicating an overlap of 10 genera between the library genera and the seed bacterial communities. Among the shared genera, *Pseudomonas*, *Stenotrophomonas*, and *Rahnella* report high relative abundance overall. *Achromobacter, Brevundimonas*, *Rhizobium* and *Variovorax* were found in all uninoculated selection types, with lower abundance; *Bacillus* was detected only in resilient-selected uninoculated lineages, while *Ensifer* and *Microbacterium* were found only in watered uninoculated lineages. *Agrobacterium* was absent from uninoculated lineages, while *Rhodococcus, Plantibacter*, and *Niallia* were not detected in any lineage types, suggesting that these genera were not present in the resident seed communities.

**Figure 4.**
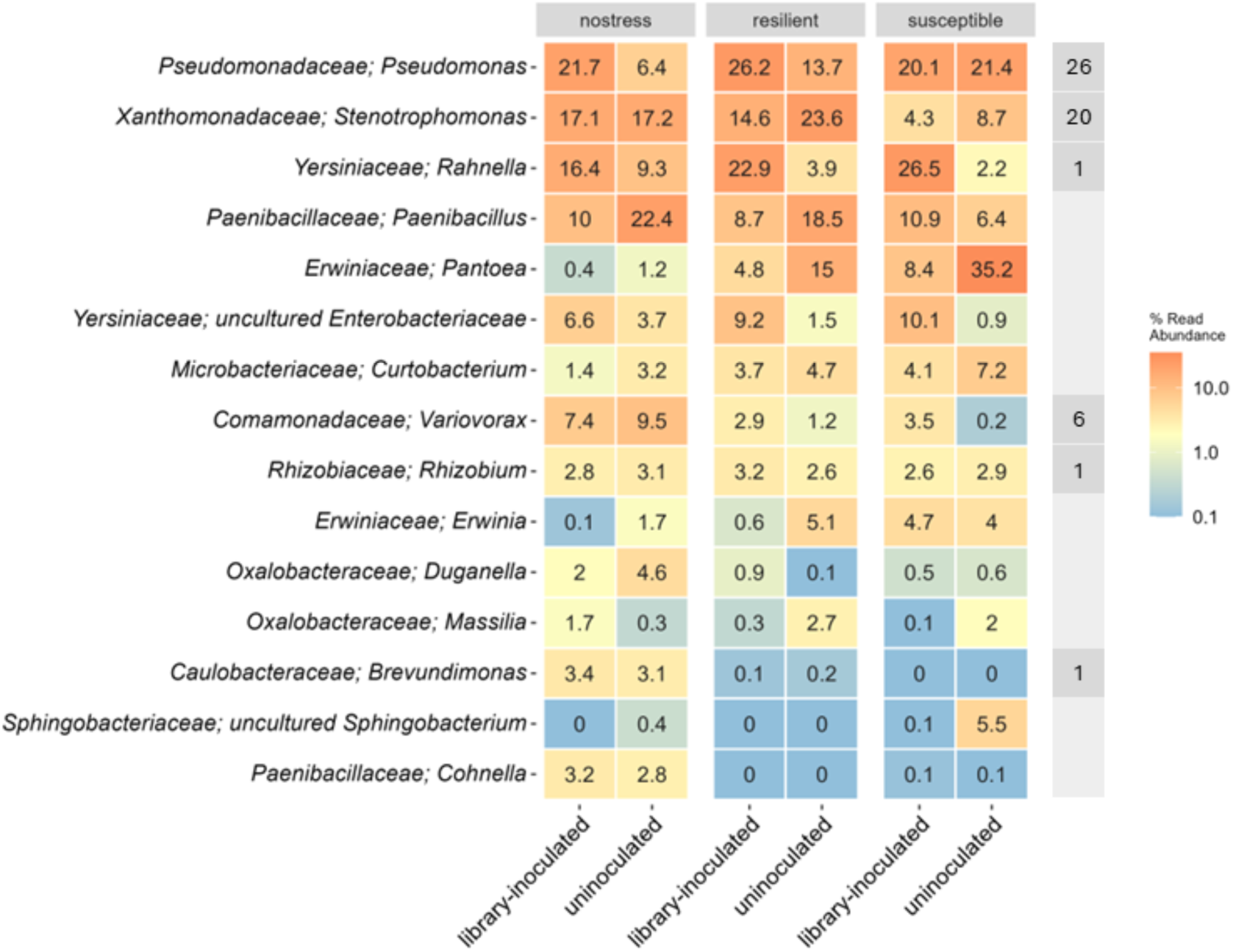
Heat map summarizes the relative abundance (as percentage of the total within each treatment) of the 15 most abundant bacterial genera in the rhizosphere of plants from each selection regime and inoculation treatment. The right-most column indicates whether each genus is included in the library by specifying the number of library strains belonging to that genus. Other library genera are not included here, as they are not represented among the 15 most abundant genera overall.

To assess the successful establishment of genera into the rhizosphere microbiome belonging to the inoculated library, we performed a differential abundance analysis focusing on the genera included in the library. A few bacterial genera represented in the inoculated library were more differentially abundant in library-inoculated compared to uninoculated treatments (Fig. 5). Specifically, *Variovorax* and *Rahnella* were significantly enriched in both resilient and susceptible inoculated lineages, and *Achromobacter* was significantly overrepresented in inoculated lineages of all selection regimes.

**Figure 5.**
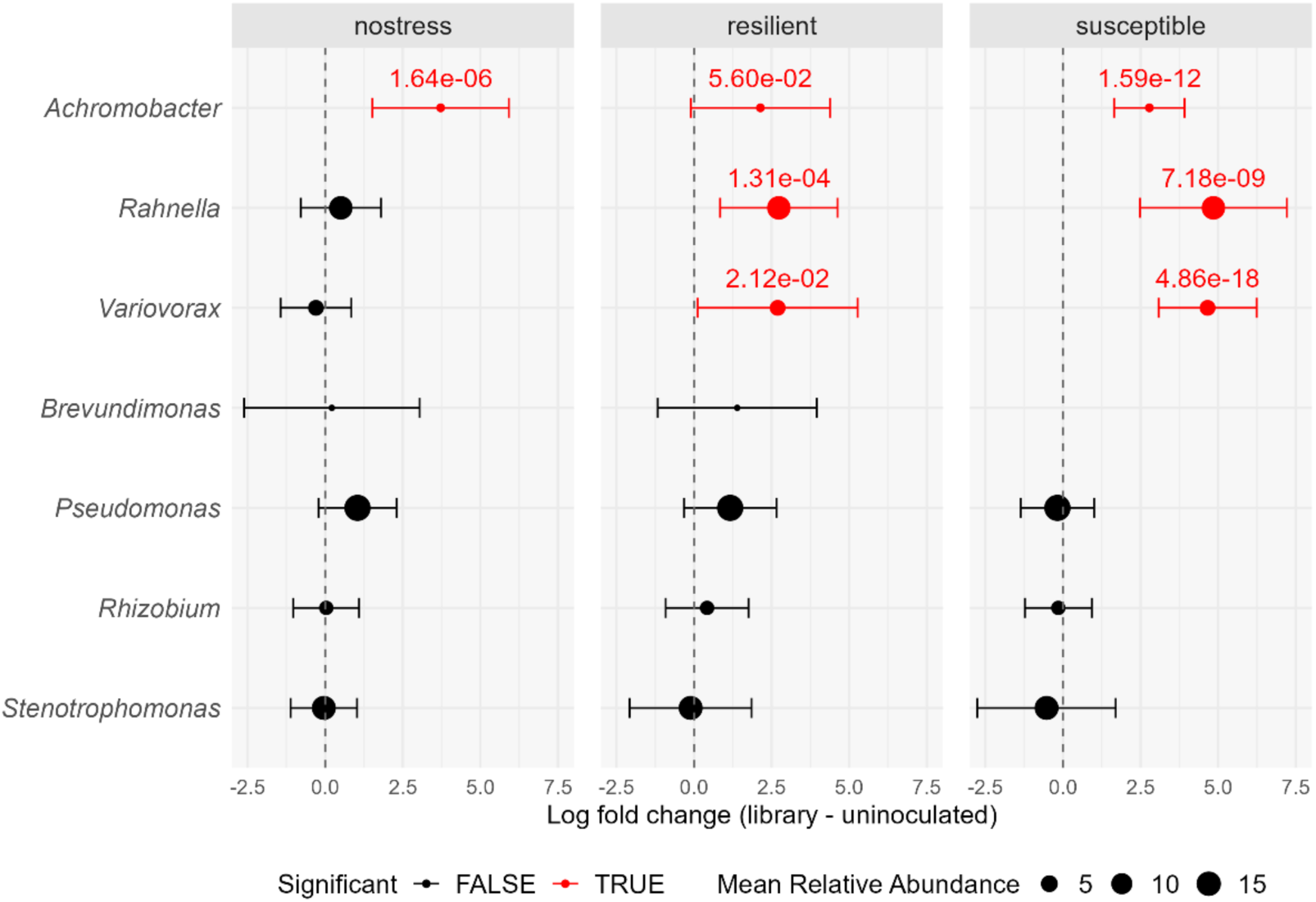
Differential abundance of bacterial genera between library-inoculated and uninoculated samples within each selection type (nostress, resilient, susceptible). Dots on the right side of the vertical line indicate enrichment in library-inoculated lineages. Data points indicate log fold changes (± 95% CI), and are sized according to the mean percent relative abundance of each genus across all treatments, to highlight dominant groups. Red points denote significantly differentially abundant genera (q-value < 0.1) for a given comparison, with q-value noted above the point. The missing comparison for *Brevundimonas* under susceptible selection is due to the especially low relative abundance of this genus in the treatments being compared. Genera with p-value = 0 for a given comparison were regarded as artifacts and excluded.

### Bacterial groups enriched in resilience-selected lineages

We investigated which microbiome members may distinguish the resilient-selected lineages, compared to the susceptible-selected lineages. In uninoculated plants, resilient-selected lineages harbored a higher differential abundance of bacterial groups including *Variovorax*, *Stenotrophomonas*, and *Rahnella*, albeit the differences were not statistically significant (Fig. 6). Instead, *Pseudomonas* was underrepresented in resilient compared to susceptible uninoculated lineages (p=0.0048, q=0.0783). In library-inoculated plants, *Stenotrophomonas* was significantly more abundant in resilient-selected plants compared to susceptible-selected plants (p=0.0015, q=0.0245), while *Erwinia*’s abundance was significantly lower (p=0.0013, q=0.0209). Other groups did not have a significantly different abundance between the two selection types. Notably, very low abundant genera may have still been significantly differentially abundant between selection types, mostly due to their absence (i.e. lack of ASV counts) in certain samples of the other treatments, therefore those are disregarded here.

**Figure 6.**
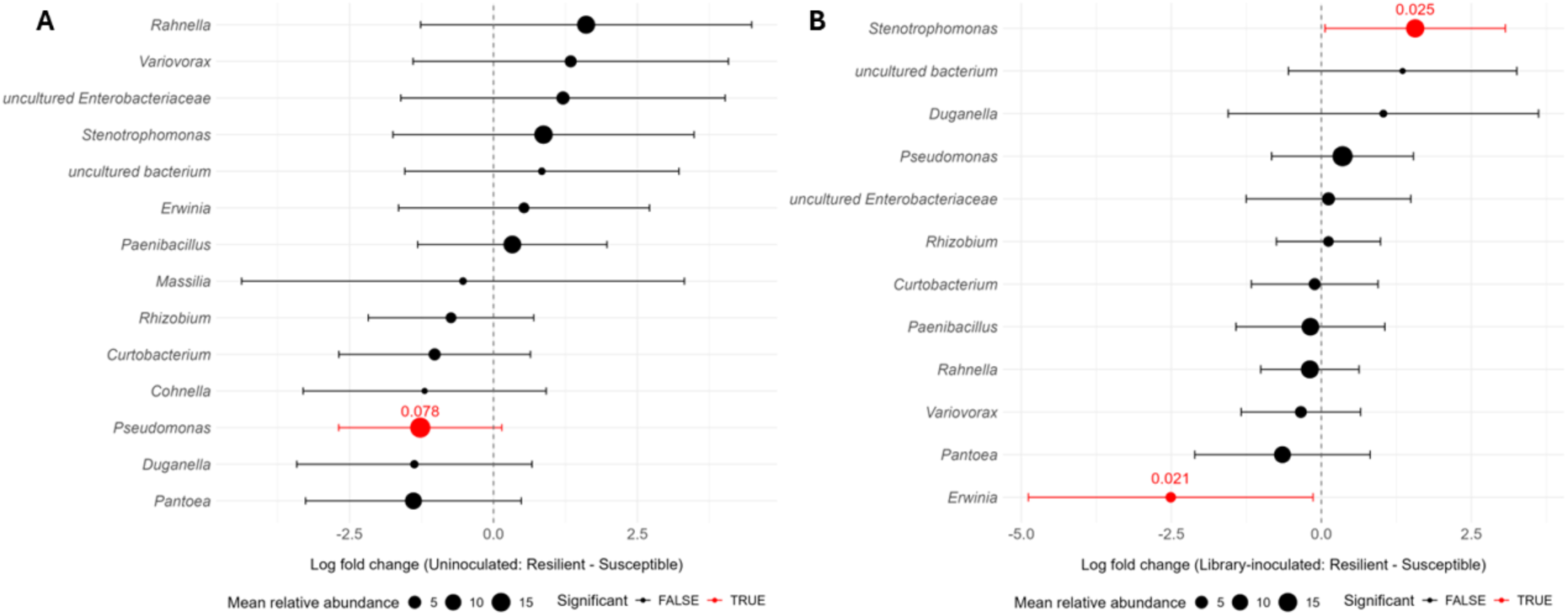
Differential abundance of bacterial genera between resilient and susceptible lineages of uninoculated (A) and library-inoculated (B) treatments. Dots on the right ride of the vertical line indicate enrichment in resilient lineages, while dots on the left side indicate enrichment in susceptible lineages. Only genera with overall relative abundance > 1% were included in the plot. Data points represent the log fold change in the abundance of each genus between the two compared treatments (± 95% CI). Notable genera with significantly higher or lower relative abundance are shown in red. Point size represents the overall relative abundance of each genus across all treatments.

### Testing *Rahnella* and *Stenotrophomonas* library strains for enhancing wheat drought resilience

We tested whether specific strains from the inoculated wheat isolate library could enhance drought resilience when inoculated into the wheat rhizosphere within one cultivation cycle. We chose *Rahnella aquatilis* strain D9 (the only strain in the library assigned to this genus), because of the significantly higher relative abundance of *Rahnella* observed in library-inoculated lineages under drought, despite its very low representation in the library. *Rahnella* also maintained higher abundance overall compared to other enriched genera. All abundant *Rahnella* ASVs from successive cultivation experiments were matched (≥99.7%) to the 16S rRNA gene sequence of strain D9. Secondly, we tested *Stenotrophomonas*, since it was the only genus which was enriched in resilient lineages, specifically under inoculation of the library. All abundant *Stenotrophomonas*-assigned ASVs found in the wheat rhizosphere after the successive cultivation experiment could be matched exactly (100%) to a set of *Stenotrophomonas* library strains (Fig. S3). All the matched *Stenotrophomonas* strains of the library, except the human-pathogenic *S. maltophilia* assigned strains, were tested in equal proportions in a combined inoculum. Plant performance was evaluated daily after the onset of drought stress using a visual scoring system (Fig. S4). A linear mixed-effects model indicated no significant effect of inoculation treatment (p = 0.73) nor a treatment × day interaction (p = 0.57) on plant scores, showing that neither inoculant improved drought performance relative to the control. Pairwise comparisons between treatments within each day also revealed no significant differences (all p > 0.15). Notably, substantial variability in plant performance was observed among replicates within each treatment.

## Discussion

We inoculated wheat seedlings with a community of 86 strains isolated from the wheat rhizosphere and rhizoplane, in order to provide them with a variety of ecologically relevant microorganisms to form interactions with. Despite surface sterilization of the seeds prior to cultivation, we expected the emergence of bacteria from the seeds into the rhizosphere under uninoculated cultivation of the wheat plants. This is in accordance with previous studies confirming the presence of bacteria embedded in the surface layers of the wheat seed capable of root colonization (Kuźniar *et al*. 2020; Garrido-Sanz and Keel 2025). Such endophytic bacteria include the genera *Erwinia*, *Paenibacillus*, *Variovorax*, *Pseudomonas*, *Stenotrophomonas*, *Flavobacterium*, and *Bacillus* (Robinson *et al*. 2016). In our study, the uninoculated treatments show signs of carry-over and accumulation of seed bacteria across cycles. However, this is also a component of microbiome development in lineages inoculated with the library. Possibly, the enhanced colonization ability of seed bacteria across the experiment also influenced their final abundance. For instance, endophytes may have a competitive advantage in colonizing the roots before other microbes from external soil (Garrido-Sanz and Keel 2025). We recognize this as a natural phenomenon, as many plants carry over endophytes inside or on their seeds to the next generation of plants (Truyens *et al*. 2015; Shade, Jacques and Barret 2017). Seed-borne microorganisms have been found to be a fundamental component of rhizosphere microbiome assembly, especially in younger plants (Johnston-Monje, Gutiérrez and Lopez-Lavalle 2021). Garrido-Sanz and Keel demonstrated that seed-borne rhizosphere bacteria accounted for a majority of the relative abundance of the wheat rhizosphere community obtained after sequential rhizosphere propagation of wheat seedlings, while only a minor fraction derived from the inoculated soil community (Garrido-Sanz and Keel 2025). Our results indicate the successful establishment of known seed bacteria, both under uninoculated conditions and with library inoculation. However, library inoculation produced different rhizosphere bacterial communities under each selection type, compared with growth in sterile soil, reaffirming the importance of the initial community composition in the active development of early rhizosphere microbiomes. Nevertheless, the genera included in the library had varying degrees of success in rhizosphere colonization. *Achromobacter* was significantly enriched in all library-inoculated lineages, albeit keeping low relative abundance overall. *Rahnella* and *Variovorax* were instead significantly enriched in both selection types under drought. *Rahnella*, and *Achromobacter* were represented by only one strain each in the 86-strain library, while six *Variovorax* isolates were present in the inoculated library. Given the high relative abundance of *Rahnella* and *Variovorax* overall in the rhizosphere samples compared to their very low representation in the library, we hypothesize that their successful establishment was driven by microbial interactions arising from library inoculation under drought conditions, and not solely by an exceptional colonization ability of the inoculated strains. Possibly, the inoculated drought treatments favored the establishment of endophytic *Variovorax* and *Rahnella* in addition to the strains added as part of the library.

However, we were unable to distinguish inoculated from resident endophytic *Rahnella* and *Variovorax* due to their high sequence similarity. *Rahnella* reached higher relative abundance in both library-inoculated lineage types under drought compared with the library-inoculated watered lineages. Interestingly, *Pseudomonas* and *Stenotrophomonas* did not have a markedly higher abundance in library-inoculated lineages as a whole, despite being abundant in the inoculated community. When subsequently testing *Rahnella* library strain D9 under drought conditions we did not observe marked positive effects on wheat drought resilience compared to the uninoculated control. *Rahnella* is a gram-negative bacterium found in soil, plant rhizospheres, animal samples, and water environments (Lindberg *et al*. 1998; Brady *et al*. 2022). *R. aquatilis* has been characterized as an opportunistic human pathogen (Matsukura *et al*. 1996; Kuzdan *et al*. 2015). *Rahnella* strains have been associated with plant disease in oak trees and onion plants (Brady *et al*. 2017; Asselin *et al*. 2019), while certain *Rahnella* species have been found to provide plant growth-promoting effects (Peng *et al*. 2019; Yuan *et al*. 2020; Li *et al*. 2021a; Cheto *et al*. 2023). To our knowledge, *Rahnella* has not been directly linked to improved drought resilience in plants. Our results indicate that this bacterium can colonize the wheat rhizosphere abundantly under drought conditions, potentially through interaction with other community members.

Differential abundance analysis comparing lineages selected for drought resilience and susceptibility, highlighted few genera with significantly different abundance. When the strain library was not inoculated, only *Pseudomonas* was significantly less abundant in resilient compared to susceptible lineages. This was not the case when the strain library was inoculated, possibly aided by the abundance of *Pseudomonas sp.* strains present in the library itself. Considering the high abundance of *Pseudomonas* across all selection types, we regard that a significant detrimental effect on drought resilience by the genus *Pseudomonas* is unlikely. Further investigation on *Pseudomonas* strains present in each lineage type is needed to clarify this. Under library-inoculated conditions, *Erwinia* was significantly less abundant in resilient-compared with susceptible-selected lineages, being almost absent from plants of both resilient and watered lineages. Instead, *Stenotrophomonas* was significantly more abundant in resilient-selected lineages compared with susceptible-selected lineages under library-inoculation. The sequences of ASVs assigned to *Stenotrophomonas* with most reads could be matched exactly (100% coverage) to the 16S rRNA gene sequence of several strains belonging to the inoculated library (Fig. S3), indicating that these strains may be of relevance to the plant, especially those reporting higher relative abundance in inoculated than uninoculated lineages. Nevertheless, when we subsequently tested a combined inoculum composed of 10 *Stenotrophomonas* library strains, no positive effects on drought resilience were observed, compared to the uninoculated control. Yet, representatives of the *Stenotrophomonas* genus have been characterized for enhancing drought tolerance in the form of improved plant growth parameters when inoculated in various crops, including chickpea (Sharma *et al*. 2023), lentils (Niza-Costa *et al*. 2022), as well as wheat (Akköprü *et al*. 2025).

Our results suggest that if *Rahnella* and *Stenotrophomonas* have the ability to enhance drought resilience, they may do so through more complex interactions involving other members of the rhizosphere community, which could not be replicated under the single inoculation of these two genera, even in presence of the resident seed community. Moreover, inoculated *Rahnella* and *Stenotrophomonas* strains may differ in their functionality from their counterparts originating from the seeds. Further characterization of seed-derived strains is needed to clarify this. Alternatively, the enrichment of the two genera in the successive cultivation experiment may be the product of plant-microbiome interactions formed under the specific conditions applied, without being directly involved in plant resilience. We suggest that these two genera may be tested further together with simplified synthetic communities (SynComs) to evaluate their colonization ability and their interaction with plants under drought stress.

Successive cultivation is carried out to highlight microbiome features which are potentially involved in the plant’s interaction with its microbiome under the different treatments. How the rhizosphere microbiome develops under a period of repeated perturbations may be consistent across plants or alternatively result in a variety of outcomes. Under each stress condition, library-inoculated replicate lineages were not significantly different from each other. We previously hypothesized that the initial inoculation of the wheat strain library would expand the range of plant-microbe interactions at the plant’s disposal and therefore may develop into more dissimilar microbiomes. Instead, the replicate lineages converged to similar microbiomes, indicating a similar response to each condition. We exclude that the library itself acted as a strong homogenizing factor within the microbiome, as the alpha diversity of inoculated lineages is comparable to the respective uninoculated lineages. Conversely, uninoculated lineages became more dissimilar, especially under non-stressed conditions and resilience selection. The higher diversity of responses across lineages suggests that plant-microbe interactions were not deterministic in these conditions, and that the microbiome developed in a manner that may be more dependent on the single selected plants driving each lineage. Since endophytes are known to interact tightly with the plant, this observation raises further questions on the role of plant-endophyte associations in stress response. Recent studies have explored various models describing microbiome dynamics and perturbations, often postulating that communities under perturbations undergo stronger shifts than healthy communities, and discussing the influence of ecological memory in community dynamics across temporal variations (Halfvarson *et al*. 2017; Gonze *et al*. 2018; Khalighi *et al*. 2022).

Plant microbiomes are known to change over time, and across plant growth stages (Dibner *et al*. 2021; Peng *et al*. 2025); therefore, the timeframe used in successive cultivation experiments is relevant. The response of plants to environmental stressors may cause gradual shifts in rhizosphere microbiome composition spanning longer timescales than the one applied in this work. Azarbad *et al*. (2022) showed that plant developmental stage and previous history of water stress in field soil were strong determinants of the wheat microbiome, and that this microbiome could shift in the span of one plant generation when exposed to a different watering treatment (Azarbad *et al*. 2022). If the microbiomes selected in our experiment were extracted and reinoculated before they became stable, they could have gone on to develop differently in the following cycle, possibly causing unpredictable effects on the plant phenotype which do not match the selection rationale they come from (Swenson, Wilson and Elias 2000). However, it is not possible to fully prevent this issue in successive cultivation experiments, due to the need for consistent growth cycles ending at a defined plant growth stage. In the future, inspection of the changes in rhizosphere microbiome composition under stress conditions is required, before choosing a suitable time frame for successive cultivation.

Previous studies employing successive cultivation focused on propagating the microbiome of plants with optimal performance (De Zutter *et al*. 2021; Mueller *et al*. 2021), and this targeted approach has been shown to produce functionally beneficial microbiomes. Jochum et al. (2019) achieved delayed onset of drought stress in wheat after consecutive microbiome cycling of drought-tolerant plants (Jochum *et al*. 2019). However, we prioritized selection of both drought-resilient and drought-susceptible phenotypes in order to compare their microbiomes and identify microbial components associated with contrasting plant responses. We were able to propagate microbiomes of drought-susceptible plants without significant accumulation of debilitating stress symptoms that would have prevented growth. In library-inoculated lineages, *Erwinia* was the only significantly enriched genus in susceptible-selected lineages compared to resilient ones. In our setup, *Erwinia* originated from the seeds as it was not present among the inoculated strains of the library. Two ASVs were the primary contributors to the abundance of *Erwinia* in susceptible-selected lineages, with high abundance variability across plants. One of them was identified as *E. persicina,* a known plant pathogen (Zhang and Nan 2014). While several *Erwinia* species are characterized as plant-pathogenic (Hsieh, Huang and Erickson 2010), *Erwinia* has been found to be a major component of the wheat seed community (Huang *et al*. 2016; Robinson *et al*. 2016). It is not straight forward whether beneficial microbes are better selected for within a good-phenotype selection than a bad-phenotype selection. In fact, we hypothesize that poorly performing plants experiencing stronger stress might engage more tightly with their microbiome to select for beneficial microbes. Therefore, selecting for poorly performing plants could result in a microbiome comprising beneficial features. If this is the case, re-inoculating the microbiome from poorly performing plants into the next cycle of plants could possibly provide a beneficial microbiome as a starting point and therefore it would be an unreliable way of identifying deleterious microbiome features under successive cultivation.

The successive cultivation experiment presented in this study did not result in observable differences in plant response to drought when comparing resilience-selected and susceptibility selected treatments (Fig. S2). Nevertheless, the two selection types resulted in significantly different microbiomes, which also differed from the watered treatment. The bacterial groups involved in producing these microbiome shifts provide an indication on microbiome traits associated with wheat plants experiencing moderate drought stress. Irrespective of whether successive cultivation is directly conducive to a functional microbiome, useful information can still be derived from comparing selection lineages and can be used in single species inocula or for designing synthetic microbial communities (Xu *et al*. 2025).

## Acknowledgement

The authors thank the members of the Bacterial Interaction and Evolution group for their help with harvesting samples after the last successive cultivation step.

## Author contributions

Adele Pioppi (Conceptualization, Formal analysis, Investigation, Methodology, Visualization, Writing – original draft), Xinming Xu (Formal analysis), Sofia I. F. Gomes (Formal analysis), Mette Nicolaisen (Funding acquisition, Resources), Ákos T. Kovács (Conceptualization, Funding acquisition, Supervision, Writing – original draft).

---

**Table S1.**
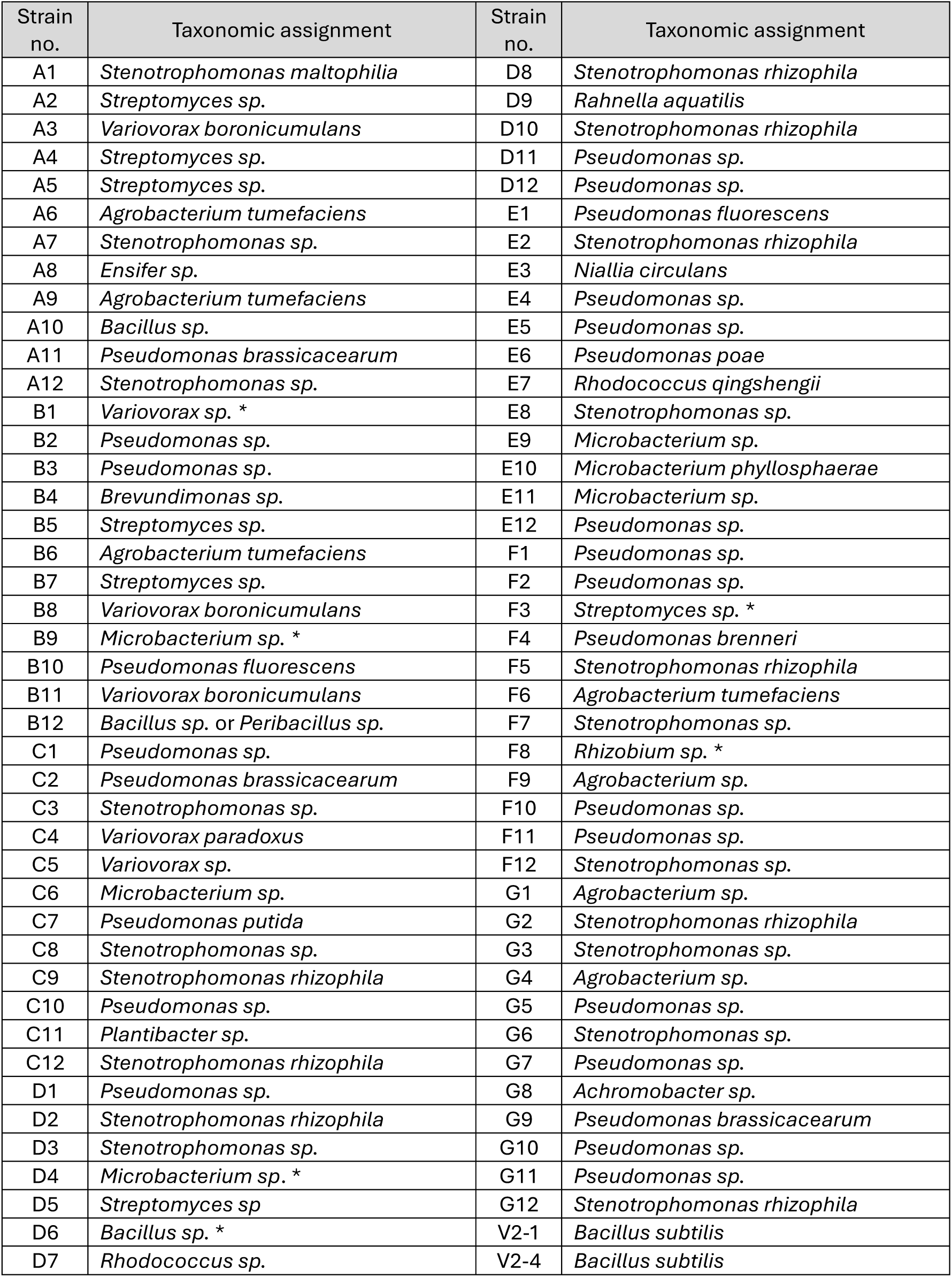
Composition of the bacterial strain library inoculated at the initiation of successive cultivation. The noted taxonomic assignment is based on sanger sequencing of the 16S rRNA gene with both forward and reverse primers. The primer sequences are included in Methods. “*” indicates that the assignment was based only on either the forward or reverse amplified fragment.

| Strain no. | Taxonomic assignment | Strain no. | Taxonomic assignment |
| --- | --- | --- | --- |
| A1 | *Stenotrophomonas maltophilia* | D8 | *Stenotrophomonas rhizophila* |
| A2 | *Streptomyces* sp. | D9 | *Rahnella aquatilis* |
| A3 | *Variovorax boronicumulans* | D10 | *Stenotrophomonas rhizophila* |
| A4 | *Streptomyces* sp. | D11 | *Pseudomonas* sp. |
| A5 | *Streptomyces* sp. | D12 | *Pseudomonas* sp. |
| A6 | *Agrobacterium tumefaciens* | E1 | *Pseudomonas fluorescens* |
| A7 | *Stenotrophomonas* sp. | E2 | *Stenotrophomonas rhizophila* |
| A8 | *Ensifer* sp. | E3 | *Niallia circulans* |
| A9 | *Agrobacterium tumefaciens* | E4 | *Pseudomonas* sp. |
| A10 | *Bacillus* sp. | E5 | *Pseudomonas* sp. |
| A11 | *Pseudomonas brassicacearum* | E6 | *Pseudomonas poae* |
| A12 | *Stenotrophomonas* sp. | E7 | *Rhodococcus qingshengii* |
| B1 | *Variovorax* sp. * | E8 | *Stenotrophomonas* sp. |
| B2 | *Pseudomonas* sp. | E9 | *Microbacterium* sp. |
| B3 | *Pseudomonas* sp. | E10 | *Microbacterium phyllosphaerae* |
| B4 | *Brevundimonas* sp. | E11 | *Microbacterium* sp. |
| B5 | *Streptomyces* sp. | E12 | *Pseudomonas* sp. |
| B6 | *Agrobacterium tumefaciens* | F1 | *Pseudomonas* sp. |
| B7 | *Streptomyces* sp. | F2 | *Pseudomonas* sp. |
| B8 | *Variovorax boronicumulans* | F3 | *Streptomyces* sp. * |
| B9 | *Microbacterium* sp. * | F4 | *Pseudomonas brenneri* |
| B10 | *Pseudomonas fluorescens* | F5 | *Stenotrophomonas rhizophila* |
| B11 | *Variovorax boronicumulans* | F6 | *Agrobacterium tumefaciens* |
| B12 | *Bacillus* sp. or *Peribacillus* sp. | F7 | *Stenotrophomonas* sp. |
| C1 | *Pseudomonas* sp. | F8 | *Rhizobium* sp. * |
| C2 | *Pseudomonas brassicacearum* | F9 | *Agrobacterium* sp. |
| C3 | *Stenotrophomonas* sp. | F10 | *Pseudomonas* sp. |
| C4 | *Variovorax paradoxus* | F11 | *Pseudomonas* sp. |
| C5 | *Variovorax* sp. | F12 | *Stenotrophomonas* sp. |
| C6 | *Microbacterium* sp. | G1 | *Agrobacterium* sp. |
| C7 | *Pseudomonas putida* | G2 | *Stenotrophomonas rhizophila* |
| C8 | *Stenotrophomonas* sp. | G3 | *Stenotrophomonas* sp. |
| C9 | *Stenotrophomonas rhizophila* | G4 | *Agrobacterium* sp. |
| C10 | *Pseudomonas* sp. | G5 | *Pseudomonas* sp. |
| C11 | *Plantibacter* sp. | G6 | *Stenotrophomonas* sp. |
| C12 | *Stenotrophomonas rhizophila* | G7 | *Pseudomonas* sp. |
| D1 | *Pseudomonas* sp. | G8 | *Achromobacter* sp. |
| D2 | *Stenotrophomonas rhizophila* | G9 | *Pseudomonas brassicacearum* |
| D3 | *Stenotrophomonas* sp. | G10 | *Pseudomonas* sp. |
| D4 | *Microbacterium* sp. * | G11 | *Pseudomonas* sp. |
| D5 | *Streptomyces* sp. | G12 | *Stenotrophomonas rhizophila* |
| D6 | *Bacillus* sp. * | V2-1 | *Bacillus subtilis* |
| D7 | *Rhodococcus* sp. | V2-4 | *Bacillus subtilis* |

**Table S2.** Results for pairwise adonis comparing each combination of treatments. * indicates significant differences (p-value=0.001, p-adjusted = 0.015).

| Comparison | Df | SumsOfSqs | F.Model | R2 | p.value | p.adjusted | sig |
| --- | --- | --- | --- | --- | --- | --- | --- |
| uninoculated_resilient vs library_nostress | 1 | 1.916057 | 12.05325 | 0.316747 | 0.001 | 0.015 | * |
| uninoculated_resilient vs uninoculated_nostress | 1 | 1.264123 | 7.37286 | 0.208433 | 0.001 | 0.015 | * |
| uninoculated_resilient vs library_resilient | 1 | 1.440572 | 7.816309 | 0.218233 | 0.001 | 0.015 | * |
| uninoculated_resilient vs library_susceptible | 1 | 1.586325 | 7.804712 | 0.21798 | 0.001 | 0.015 | * |
| uninoculated_resilient vs uninoculated_susceptible | 1 | 1.000765 | 4.433318 | 0.13669 | 0.001 | 0.015 | * |
| library_nostress vs uninoculated_nostress | 1 | 0.92671 | 8.011352 | 0.235549 | 0.001 | 0.015 | * |
| library_nostress vs library_resilient | 1 | 0.796253 | 6.148207 | 0.191246 | 0.001 | 0.015 | * |
| library_nostress vs library_susceptible | 1 | 1.375052 | 9.172123 | 0.260778 | 0.001 | 0.015 | * |
| library_nostress vs uninoculated_susceptible | 1 | 3.169749 | 18.20325 | 0.411808 | 0.001 | 0.015 | * |
| uninoculated_nostress vs library_resilient | 1 | 1.533839 | 10.64397 | 0.275437 | 0.001 | 0.015 | * |
| uninoculated_nostress vs library_susceptible | 1 | 1.933876 | 11.86042 | 0.297549 | 0.001 | 0.015 | * |
| uninoculated_nostress vs uninoculated_susceptible | 1 | 2.903914 | 15.65133 | 0.358553 | 0.001 | 0.015 | * |
| library_resilient vs library_susceptible | 1 | 0.421312 | 2.395179 | 0.078801 | 0.011 | 0.165 |  |
| library_resilient vs uninoculated_susceptible | 1 | 1.928653 | 9.721768 | 0.257723 | 0.001 | 0.015 | * |
| library_susceptible vs uninoculated_susceptible | 1 | 1.887102 | 8.682963 | 0.236703 | 0.001 | 0.015 | * |

**Figure S1.**
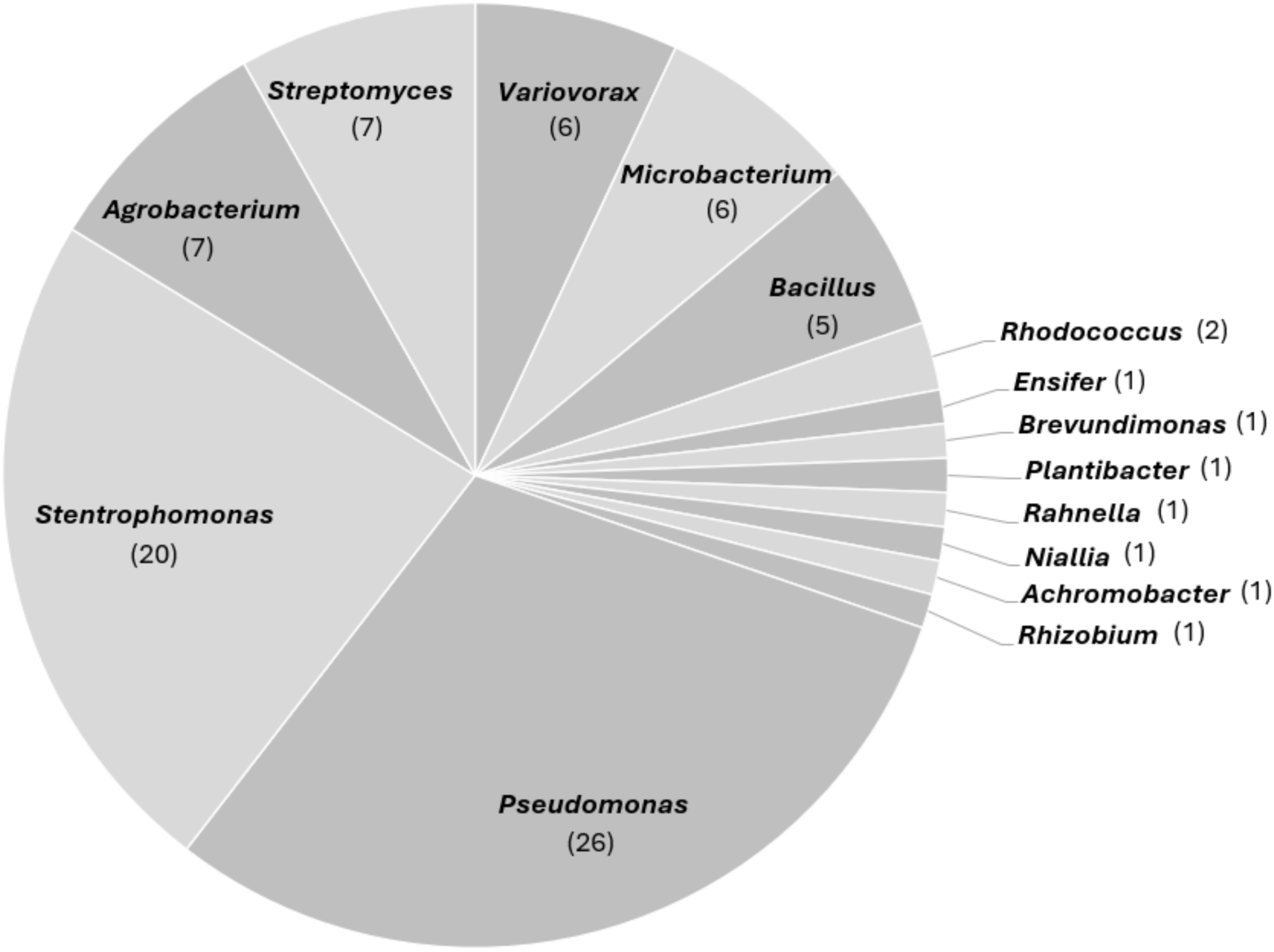
Distribution of bacterial genera included in the library, with the number of strains belonging to each genus noted in parentheses.

**Figure S2.**
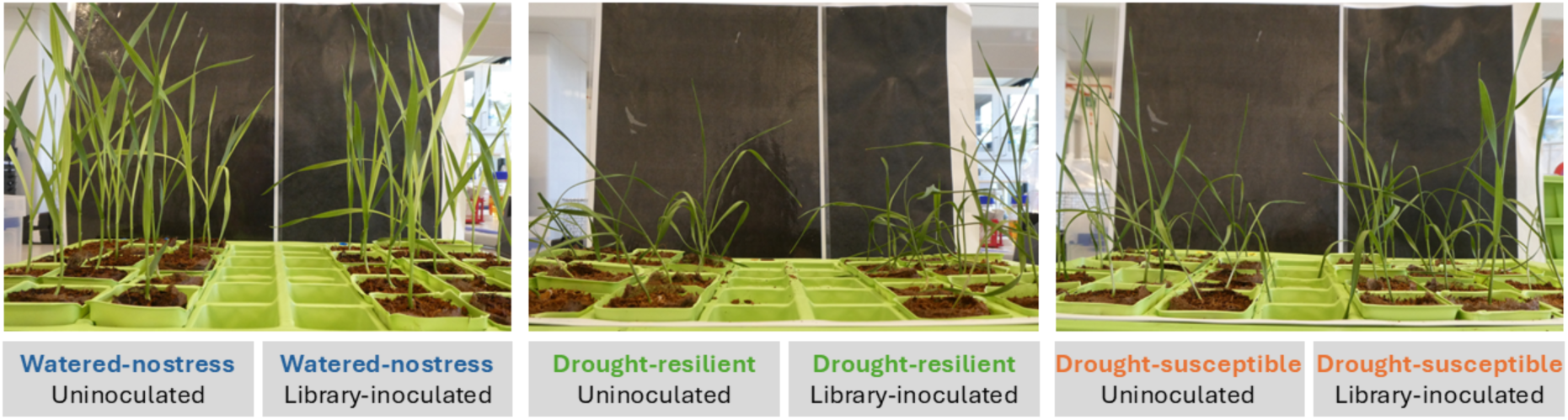
Photos of plants taken on the last day of the 4th cycle, before starting rhizosphere extraction. Selection types and treatments are noted below, distinguishing Uninoculated and Library-inoculated lineages of each selection type.

**Figure S3.**
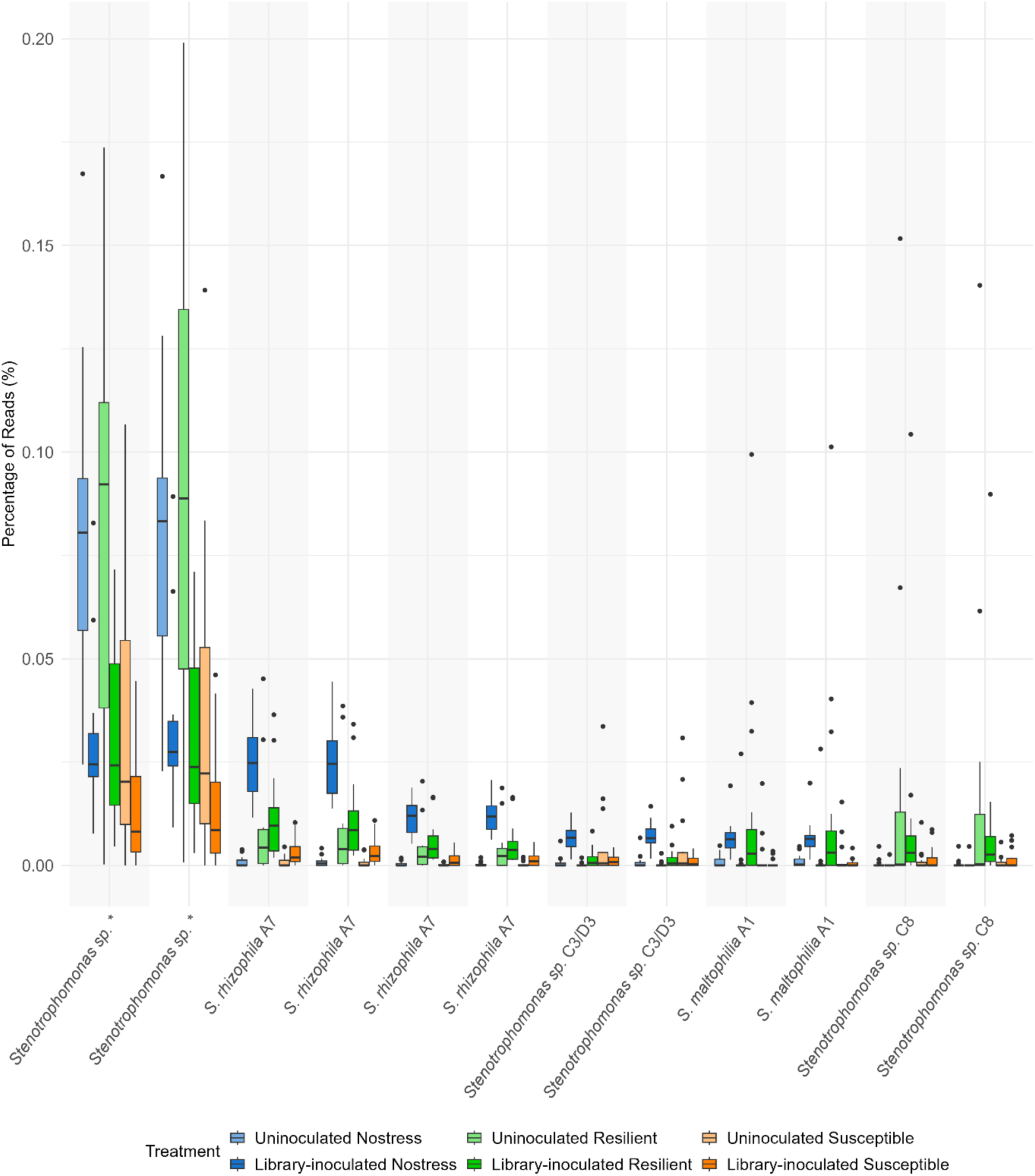
Relative abundance (as percentage of reads) of 12 ASVs assigned to *Stenotrophomonas* within each treatment. Labels indicate which *Stenotrophomonas* library strain each ASV could be matched to, with 100% coverage of the V3-V4 region of the 16S gene. The asterisk (*) indicates an exact match of two ASVs to multiple library strains, specifically C12, D2, D8, D10, E2, F5, F12. The ASVs and corresponding matched library strains are ordered according to similarity in their distribution across treatments.

**Figure S4.**
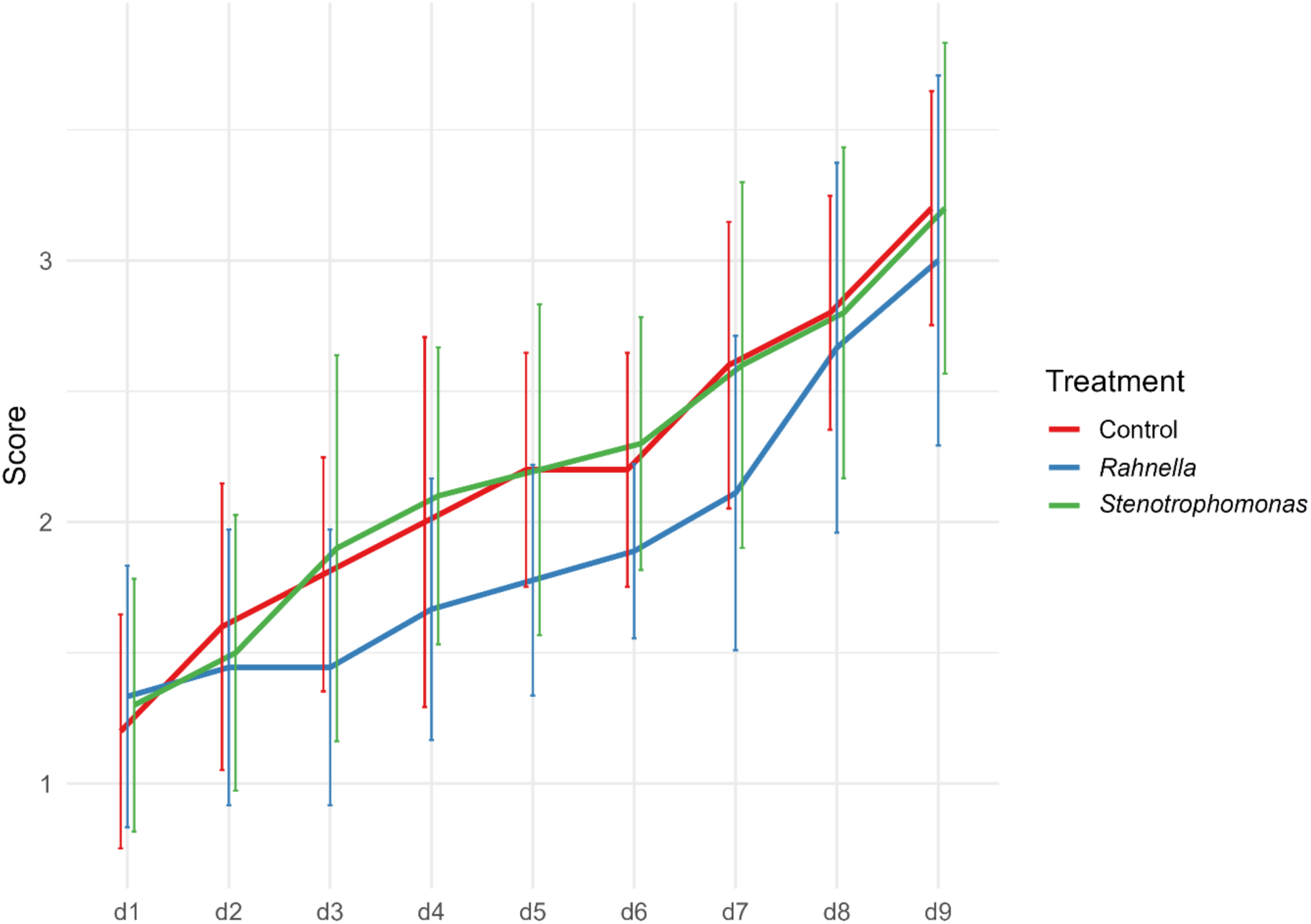
Performance scores of wheat plants under drought based on a daily visual assessment. The assessment started on the first day where any sign of drought stress were visible (d1). Days are indicated at the bottom of the plot. The main line corresponds to the mean score on each day, and error bars indicate standard deviation (SD). A slight horizontal shift is added to each treatment line to prevent overlapping of error bars.

